# Multiscale temporal tuning of force generation complex machinery governs cortical microtubule interactions during the first mitotic division in *C. elegans*

**DOI:** 10.64898/2026.01.16.699965

**Authors:** G. Alan Edwards, John B. Linehan, Amy Shaub Maddox, Paul S. Maddox

## Abstract

Dynein is an essential microtubule motor whose many roles in mitosis complicate efforts to resolve its spatiotemporally dynamic regulation. We previously established a method to classify single particles of the conserved cortical force generation complex (dynein—LIN-5^NuMA^—GPR-1/2^LGN^—Gα^Gαi^), as free or interacting with microtubules. Here, we report the results of depleting force generation complex components and regulators. Depleting LIS-1 reversed force asymmetry, while depletion of the regulatory subunit SUR-6^PP2A-B55^ significantly increased incidence of dynein trajectories with microtubule-interacting behavior during prophase. We next applied our classification scheme to the dynein anchor LIN-5. Microtubule-interacting LIN-5 trajectories were posteriorly enriched during anaphase, consistent with our dynein data and prior fluorescence studies. SUR-6^PP2A-B55^ depletion did not alter LIN-5 kinetics, suggesting regulation of a dynein specific function rather than regulation via LIN-5. We found evidence that two distinct regulators of force generation complex behavior, microtubule interactions and cortical flows, govern on the millisecond and second time scales, respectively. Our observations provide novel insight into the regulation of cortical dynein and the coupling among the cell membrane, actomyosin cortex, force generation complex, and microtubules that position the mitotic spindle.

## Introduction

During embryogenesis and throughout development, asymmetric cell division is a major mechanism for generating cell diversity. The position of the mitotic spindle during mitosis sets the division plane and helps determine the size and fate of the daughter cells. Dynamic microtubule polymers emanate from two centrosomes and interact with dynein anchored at the cortex, which generates forces that position the spindle and therefore the cleavage plane. In *C. elegans*, asymmetric division of the zygote is required for normal development (Jankele *et al*., 2021).

The dimeric cytoplasmic dynein complex (hereafter referred to as dynein) is a minus-end directed microtubule motor powered via ATP hydrolysis to drive processive motion along microtubules. Although dynein is best known for cargo transport along the microtubule lattice, it can also generate force on a microtubule by cross-linking and bundling adjacent microtubules, or by tethering to a membrane and interacting with microtubules (Laan *et al*., 2012a; Roberts *et al*., 2013; Mazel *et al*., 2014).

During cell division, dynein plays multiple roles in establishing and maintaining the shape, size, and position of the mitotic spindle (Merdes *et al*., 1996; Gönczy *et al*., 1999). Dynein tethered to the cortex pulls on microtubules emanating from centrosomes, dynamically positioning the spindle (Gusnowski and Srayko, 2011; Laan *et al*., 2012b), while dynein at the kinetochore mediates the kinetochore-microtubule attachment (Yang *et al*., 2007; Vorozhko *et al*., 2008; Barisic *et al*., 2014). Dynein anchored at the nuclear envelope drives nuclear envelope breakdown and spindle assembly (Salina *et al*., 2002). In the *C. elegans* zygote, dynein anchored to the pronuclear envelope and cortex interacts with centrosome-associated microtubules to achieve centrosome separation and nuclear positioning prior to nuclear envelope breakdown (Gönczy *et al*., 1999; Robinson *et al*., 1999; Crisp *et al*., 2006; Splinter *et al*., 2010; De Simone *et al*., 2016, 2018; De Simone and Gönczy, 2017).

At the cortex, dynein is tethered by the conserved tripartite complex consisting of GOA-1/2—GPR-1/2—LIN-5 (Gα—LGN—NuMA in humans, reviewed in (Kiyomitsu, 2019)). Asymmetric force generation arises from force generators’ unequal cortical distribution, mediated by polarity cues (Rodriguez-Garcia *et al*., 2018). Time-resolved fluorescence imaging confirmed the organization and necessity of these proteins and indicated that cortical polarization of GPR-1/2 is initially driven by actomyosin flows; GPR-1/2 in turn recruits LIN-5, which then tethers dynein to the cortex (De Simone *et al*., 2016; Fielmich *et al*., 2018; Okumura *et al*., 2018). Consequently, cortically bound dynein is distributed unequally along the anterior–posterior axis, generating the force imbalance required for posteriorized spindle positioning and asymmetric cell division.

Several studies have explored the architecture and behavior of the force generation complex. Single particle studies of cortical dynein revealed two populations – one dependent on LIN-5, and the other associated with microtubule end-binding proteins (Schmidt *et al*., 2017; Rodriguez-Garcia *et al*., 2018). Dynein particles exhibiting diffusive-like (as opposed to directed) trajectories were asymmetrically distributed and dependent on LIN-5 and GPR-1/2. While informative, these studies lacked the temporal resolution needed to describe dynein motion during active force generation. Live-cell imaging experiments investigating other components of the force generation complex have used proxies or fluorescence intensity measurements (Park and Rose, 2008; Fielmich *et al*., 2018). The spatial resolution of these experiments is not suited for exploring kinetic relationships and transient interactions. Thus, how dynein and the force generation complex dynamically interact to generate force on microtubules and how this process is locally regulated have not been explored in great detail.

To overcome this limitation, we performed light microscopy and single-particle tracking with nanometer-scale precision and millisecond temporal resolution, to reveal variations in particle motion at biologically relevant timescales. Previously, we combined single particle tracking with total internal reflection fluorescence (TIRF) microscopy with a quantitative model to classify cortical dynein force generation behavior in the *C. elegans* zygote (see Materials and Methods; (Linehan *et al*., 2023)). In brief, the model categorized dynein trajectories according to their step-length distributions, identifying cortically tethered dynein molecules whose displacement changes were consistent with microtubule interaction while remaining anchored to the force generation complex. This approach reproduced known force asymmetries during pronuclear migration and anaphase and was sensitive to the reduction of microtubules (Park and Rose, 2008; Fielmich *et al*., 2018; Linehan *et al*., 2023).

Here, we extended that framework to investigate how cortical dynein is regulated, and how the force generation complex functions as an integrated unit. Through this analysis, we identified a previously unrecognized role for the conserved protein phosphatase 2A (PP2A B55/SUR-6) in cortical force generation. RNA-mediated depletion (RNAi) of the regulatory subunit SUR-6 hyperactivated dynein, producing a global increase in cortical force generation. We also analyzed single particle trajectory data from the dynein anchoring protein LIN-5 to gain additional insight into the relationship between dynein and the force generation complex. Two distinct temporal regimes of regulation emerged: cortical tethering of dynein and LIN-5 occurs on the millisecond scale, while LIN-5 is displaced on the second scale, similar to flows of the actomyosin cortex. Together, these results reveal that cortical dynein activity and the distribution of LIN-5 are tuned on different timescales.

## Materials and Methods

### C. elegans culture, RNAi, and microscopy

The *C. elegans* strains SV1803 (he264[eGFP::dhc-1]) and LP585 (cp288[lin-5::mNG-C1^3xFlag]) II were maintained at 20°C using standard procedures (Schmidt *et al*., 2017; Heppert *et al*., 2018). Bacterial strains containing a vector expressing double-stranded RNA under the isopropyl β-d-1-thiogalactopyranoside-inducible promoter were obtained from the Ahringer library (Bob Goldstein’s laboratory, University of North Carolina at Chapel Hill, Chapel Hill, NC) (Fraser *et al*., 2000; Kamath *et al*., 2001). Targets were confirmed by sequencing.

Adult animals in the fourth larval stage (L4) were fed bacteria harboring the empty RNAi vector (L4440) as a negative control for 24 hours before imaging or with bacteria expressing dsRNA targeting *tba-2* for 16 hours (longer TBA-2 depletion caused gross defects in oogenesis and zygote mitosis), and 24 hours for all other depletions. Embryos were dissected from treated hermaphrodites and mounted in egg buffer between a #1.5 coverslip and a 4% agarose pad and sealed with VaLaP. Embryos were imaged in prophase or anaphase at anterior or posterior poles on a Nikon TIRF microscope with a 100× Apo TIRF oil-immersion objective (NA = 1.49, Nikon), an Andor iXon3 EMCCD (SV1803) or Photometrics Prime 95B SCMOS (LP585) and NIS-Elements (Nikon) at 22°C. The incident angle of the laser was maximized to illuminate only the cortex of the embryo. Images were acquired at 40 Hz.

### Image processing and analysis

In both *C. elegans* strains, SV1803 and LP585, a high density of particles was observed at the cortex that confounded the identification of individual particles and trajectories. To account for the poor signal/noise ratio in SV1803, we photobleached the specimens for 1000 time points (27 seconds). The photobleached frames were removed before processing. To account for the high density but better signal/noise ratio in LP585, we employed a custom difference of gaussian script (Fig. S2, Movie S2). A histogram-matching photobleaching algorithm available in Fiji was applied to decrease the variability of particle intensity and tracking segmentation over time for all images (Schindelin *et al*., 2012; Miura, 2020). All image analysis was done using the TrackMate plugin in Fiji (Tinevez *et al*., 2017; Ershov *et al*., 2021, 2022). For SV1803, tracking was performed using a difference of Gaussian detection segmenter (estimated object diameter = 0.4 μm, quality threshold = 6) and simple linear assignment problem tracking (linking max distance = 0.5 μm, gap-closing distance = 0.5 μm, gap-closing max frame gap = 1). Tracks with fewer than 5 spots were discarded. Images with fewer than 50 total tracks were discarded. For LP585, tracking was performed using a difference of Gaussian detection segmenter (estimated object diameter = 0.35 μm, quality threshold = ∼0.2-0.3) and linear assignment problem tracking (frame to frame linking = 0.3 μm). Spots within 0.1 μm of the edge of the image were discarded. Tracks with fewer than 5 spots were discarded. Custom scripts are available upon request.

### Image normalization to compare SV1803 and LP585

LIN-5::mNG images were taken with a Prime 95b sCMOS camera, which allowed us to improve our field of view without sacrificing speed while improving signal-to-noise resolution with minimal loss in resolution (∼0.06 μm/pixel (EMCCD), ∼0.08 μm/pixel (sCMOS)). To compare total dynein trajectories with total LIN-5 trajectories, we divided the number of trajectories by the area contained by the trajectories for each movie. Imaging speed (∼40 Hz) and total time (∼ 1 min) were the same amongst conditions.

### Model-based trajectory classification scheme

Classifying cortical dynein as free (non-interacting) or interacting (microtubule-engaged) was performed as previously reported (Linehan *et al*., 2023). Briefly, the force generating complex was considered as a ball joint swiveling about a fixed base. Our scheme used the spherical position coordinates ρ (radial coordinate), θ (polar angle), and φ (azimuthal angle) to model the position of a fluorophore in three-dimensional space. φ was modeled as a uniform distribution because dynein trajectory turn angles were unbiased. We used step-length displacements obtained from single particle tracking data to simulate polar angles of deflection over different lengths (ρ). We performed Monte Carlo simulations to create step-length probability distributions for each maximum polar angle of deflection for free swivel (*θ*^*fmax*^) and interacting (*θ*^*imax*^) groups for each ρ. We compared the simulated probability distributions to experimental probability distributions, finding the combination of ρ, *θ*^*fmax*^, *θ*^*imax*^ that best fit experimental data (sum of squared error). The same pipeline was used to generate ρ, *θ*^*fmax*^, *θ*^*imax*^for LIN-5::mNG trajectories.

### Estimation of force generation complex length

GOA-1, GPR1/2, and LIN-5 lengths were calculated from *C. elegans* predicted structures using AlphaFold and amino acid sequences from Uniprot (Jumper *et al*., 2021; Varadi *et al*., 2022). Protein length was estimated using the greatest linear distance between polar amino acids. The file IDs accessed for this study were UPID: P45970 (LIN-5), UPID: P51875 (GOA-1), UPID: Q95QJ7 (GPR-1), UPID: Q03569 (GPR-2).

### Particle Image Velocimetry

To measure cortical flow, we downsampled our images, reducing frames by a factor of 20 to reduce computational expense. The resulting image sequence was imported into Matlab (v24.2), where the tool PIVlab (v3.08) was implemented (Thielicke and Sonntag, 2021; Thielicke, 2022). Areas were drawn in the anterior and posterior embryo to measure average flow magnitude for embryos in which both regions were clearly visible (n=8).

## Results

### Analyzing trajectories for biological interpretation: Activity and collective efficiency index

We recently described a quantitative framework to classify cortical dynein behavior using single particle trajectory data and mathematical modeling (Linehan *et al*., 2023). This approach distinguished cortically bound dynein trajectories engaged in force generation (interacting swivel) from those that were not (free swivel). We compared the total interacting trajectories across experimental conditions to verify the model against previously established benchmarks. Our model recapitulated the characteristic force asymmetry along the anterior–posterior axis in prophase and anaphase, and was sensitive to microtubule perturbation via TBA-2^β-tubulin^ depletion.

When we applied our approach to more complex perturbations, however, we identified that the number of interacting trajectories alone did not fully capture changes to the dynamics of the cortical dynein landscape. Variations in the total number of trajectories among RNAi treatments obscured differences in interacting trajectories. To compare our results between treatments, we quantified (1) the proportion of interacting dynein trajectories and (2) the magnitude of interacting trajectory with respect to the proportion. To compare proportions between treatments, we calculated the fraction of the total trajectories *N* that were interacting (*I*): “Activity” (*A*).

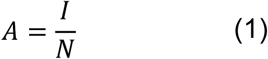

The Activity ratio did not account for the magnitude (number) of interacting trajectories. To address this limitation, we introduced a second metric, the collective efficiency index (CEI), which combines the magnitude of interacting trajectories with overall Activity. Specifically, we multiplied *I* by *A* and applied a *log*_2_ transform for symmetry and statistical analysis,

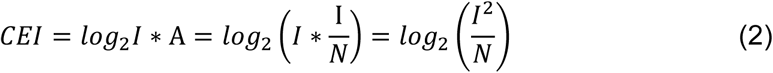

CEI therefore reflects both the absolute contributions of interacting dynein trajectories and relative proportions of populations within each condition. Finally, we quantified anterior–posterior trajectory asymmetry by comparing CEI values between embryo halves,

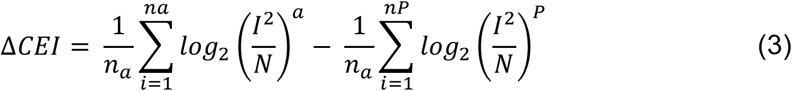

where *n* represents the number of embryos and *a*, *p* denote anterior and posterior, respectively. ΔCEI quantifies the degree to which cortical force imbalance is altered between the two embryo halves (see Fig. S1 for *in silico* demonstration of the complementarity between “Activity” and “CEI” metrics).

To investigate how cortical dynein is regulated, we applied this analytical framework to *C. elegans* embryos imaged by TIRF microscopy (Movie S1). First, we focused on prophase, when cortical pulling forces are strongest in anterior and centrosome migration and centration occur. Using RNAi, we perturbed components that influence microtubule stability (EFA-6 and TBA-2), known dynein-interacting cofactors (LIS-1 and LIN-5), and the regulatory subunit of the protein phosphatase PP2A-B55 (SUR-6), whose role in cortical dynein regulation remains unclear.

### Microtubule stability establishes baseline Activity and CEI dynamics

We re-analyzed our previously published *tba-2(RNAi)* (16h) dataset, with fewer interacting trajectories in both anterior and posterior cortices during prophase compared to control (Linehan *et al*., 2023). We tested whether the swivel model and the Activity and CEI metrics remained sensitive to changes in microtubule abundance. To create a condition in which cortical microtubules are more numerous and stable than in controls, we depleted the microtubule growth limiting factor EFA-6 (O’Rourke *et al*., 2010).

Reducing microtubule abundance by depleting TBA-2^tubulin^ significantly decreased Activity (Brown–Forsythe ANOVA, Welch’s t-test, p < 0.0001) and reduced CEI in both embryo halves (one-way ANOVA, unpaired t-test, anterior p = 0.0025; posterior p = 0.0025), while preserving the anterior–posterior asymmetry (Fig. 1 A-D). In contrast, *efa-6(RNAi)* embryos showed no change in Activity but exhibited increased CEI, particularly in the posterior (Fig. 1B-C, one-way ANOVA, unpaired t-test, anterior p = 0.0307; posterior p < 0.0001). EFA-6 depleted embryos also displayed symmetric ΔCEI, indicating a disruption of anterior–posterior force imbalance (Fig. 1D). Together, these perturbations established baseline conditions for interpreting changes in Activity and CEI and confirmed that both metrics reliably capture the impact of microtubule stability on cortical dynein-microtubule interactions.

**Figure 1.**
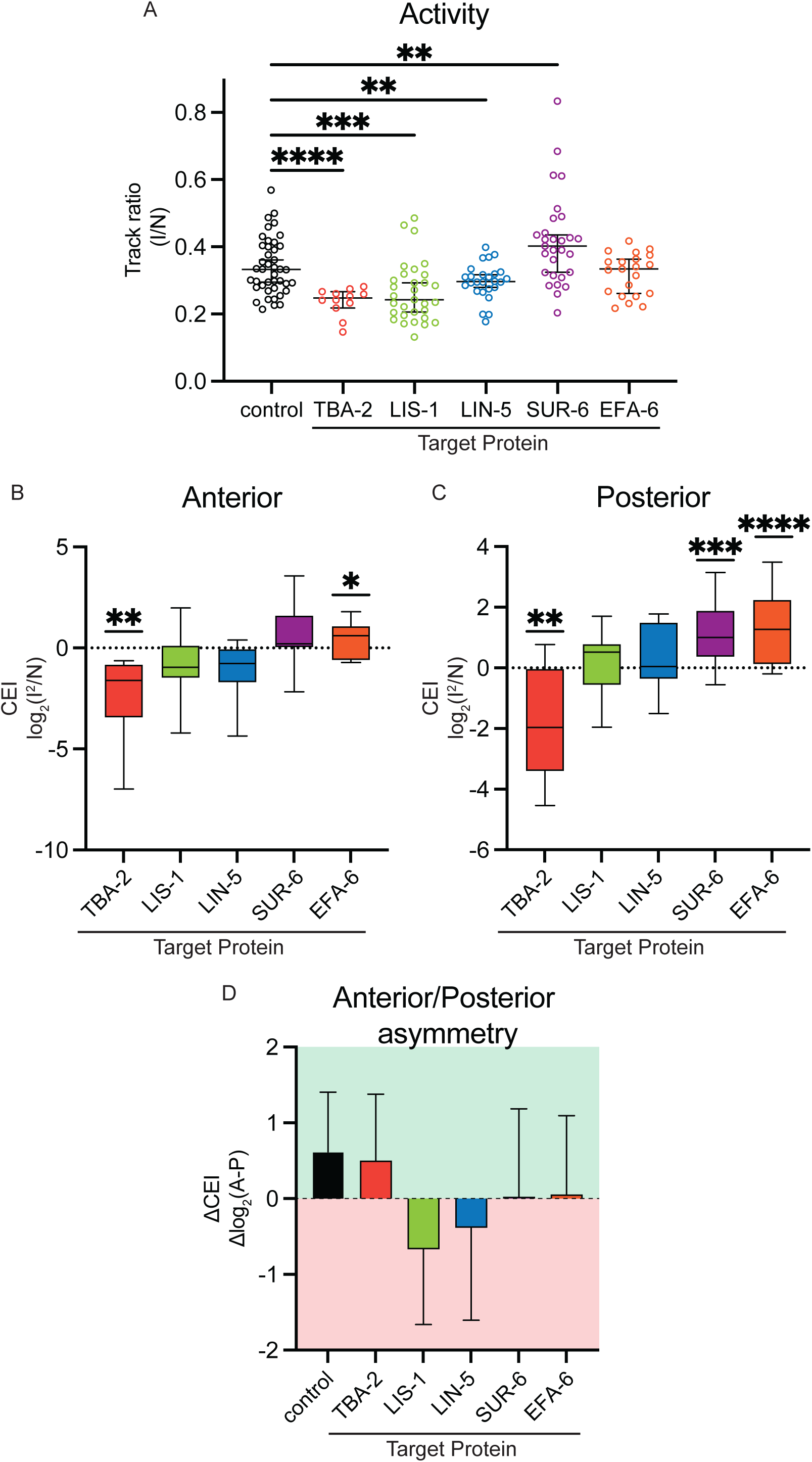
Dynein regulation revealed through swivel model Activity and collective efficiency index (CEI) metrics. (A) Activity metrics obtained from swivel data. Circles represent individual embryos. Black lines represent median ± 95% CI. (B–C) Baseline-subtracted box and whisker plots of CEI in the anterior (B) and posterior (C). Black lines represent 5th and 95th percentile. Statistics: Activity - Brown–Forsythe Anova and Welch’s t-test. CEI – one-way ANOVA and unpaired t-test. All t-tests for significance correspond to *p<0.05, **p<0.01, ***p<0.001, ****p<0.0001. (D) Anterior–posterior asymmetry measured as ΔCEI (anterior – posterior).

### LIN-5 is essential for force polarization

The anterior-posterior polarization of the dynein anchoring protein LIN-5 (downstream of GPR-1/2 and the PARs) is a well-established determinant of asymmetric force generation during cell division (Goldstein and Macara, 2007). Membrane anchoring of LIN-5 is essential for dynein-mediated cortical pulling forces (Fielmich *et al*., 2018; Jankele *et al*., 2021).

Consistent with this precedent, depletion of LIN-5 caused a significant decrease in dynein Activity (Fig. 1A, Brown–Forsythe ANOVA and Welch’s t-test, p = 0.0099). CEI decreased in the anterior and increased in the posterior cortex, eliminating the typical anterior–posterior asymmetry (Fig. 1B-D). These findings confirm that LIN-5 is essential for cortical force polarization and demonstrate that our Activity and CEI analyses accurately capture changes in force generation complex distribution and cortical dynein-microtubule interactions.

### LIS-1 activation modulates dynein force asymmetry

LIS-1 is a conserved dynein activating adaptor that has been extensively studied for its role in human lissencephaly. *In vitro*, LIS-1 enhances dynein processivity along the microtubule lattice, increasing run length and velocity, and is thought to release dynein from autoinhibition by acting as a molecular wedge (Huang *et al*., 2012; Zou *et al*., 2014). *In vivo* in *C. elegans,* LIS-1 localizes to multiple subcellular compartments, including the nuclear envelope and the cortex, but its specific contribution to cortical force generation remains unclear.

We hypothesized that if LIS-1’s sole cortical role is to activate dynein, LIS-1 depletion would reduce both Activity and CEI, but not alter the spatial distribution of dynein. Consistent with this prediction, dynein Activity was significantly decreased in *lis-1(RNAi)* embryos compared to control (Fig. 1A, Brown–Forsythe ANOVA and Welch’s t-test, p = 0.0004). CEI was slightly reduced in the anterior and modestly increased in the posterior; thus the typical anterior–posterior force asymmetry was reversed. (Fig. 1B-D). Together, these results suggest that LIS-1 positively regulates cortical dynein engagement, consistent with its role in dynein activation, and implies that LIS-1 also influences the asymmetric distribution of active dynein at the cortex.

### PP2A-B55/SUR-6 depletion hyperactivates cortical dynein

We previously identified SUR-6 as a regulator of microtubule interactions at the cortex. Embryos depleted of SUR-6 displayed centrosome separation and migration defects, which were partially attributed to SUR-6-dependent nuclear size regulation (Boudreau *et al*., 2019). Cortical microtubules polymerized with reduced velocity compared with controls, suggesting that SUR-6 positively regulates the force generation complex. This led us to ask whether SUR-6 regulates cortical force generation through dynein.

SUR-6 depletion significantly increased dynein Activity (Fig. 1A; Brown–Forsythe ANOVA, Welch’s t-test p = 0.0094). CEI was modestly elevated in the anterior but strongly increased in the posterior (Fig. 1 B–C; one-way ANOVA, unpaired t-test; p = 0.0001), while anterior–posterior asymmetry was maintained (Fig. 1D). Thus, reduction of SUR-6 raises cortical dynein activity. Although SUR-6 also functions at the nuclear envelope and centrosomes, these findings provide the first direct evidence that SUR-6 regulates (limits) cortical dynein-microtubule interactions in the *C. elegans* embryo.

### Modeling LIN-5 swivel behavior reveals the architecture of the cortical force generation complex

We considered that the relatively short trajectory lengths of dynein may not fully capture the motion of the entire force generation complex. Given that LIN-5 anchors dynein to the cortex, we reasoned that analyzing LIN-5 trajectories using the same swivel model framework would reveal complementary insights into the behavior of the force generation complex.

To visualize LIN-5, we imaged *C. elegans* embryos expressing LIN-5::mNG (Fig. 2A, Movie S3; (Heppert *et al*., 2018)) using TIRF microscopy during anaphase, when cortical forces are greatest in the anterior cortex. Step-length distribution for LIN-5 trajectories were shifted toward shorter displacements compared to dynein, with a peak step-length probability at 0.020 µm for LIN-5 in the anterior and posterior and 0.045 µm and 0.035 µm for dynein in the anterior and posterior, respectively. (Fig. 2B).

**Figure 2.**
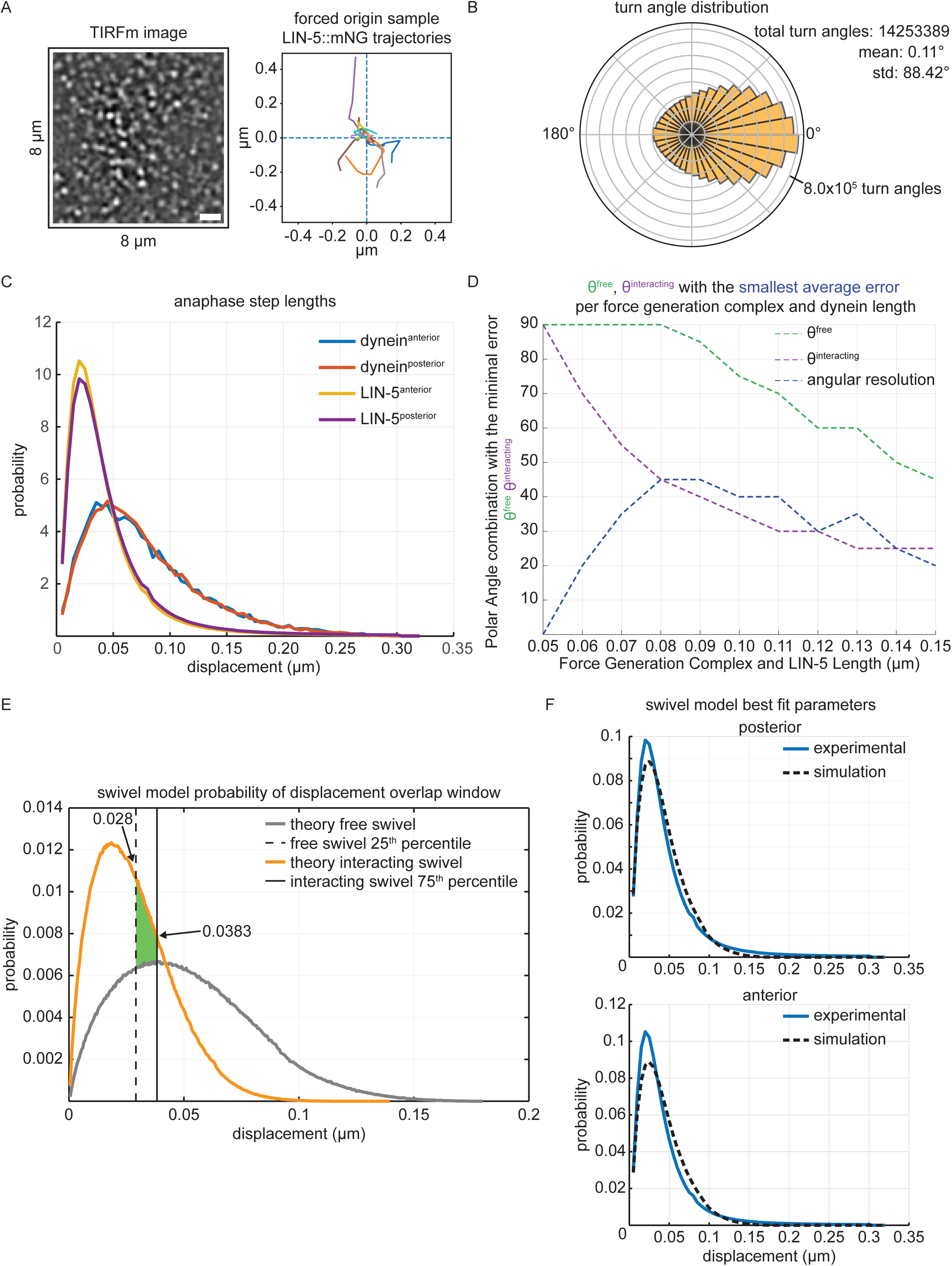
LIN-5 swivel classification scheme. (A) Representative TIRF image of lin-5::mNG cropped to 8 x 8 μm, with sample trajectories starting at the origin (0,0). x,y coordinates are in μ m. (B) LIN-5 trajectory turn angle distribution, including total number of trajectories analyzed across all control embryos, with mean and standard deviation (std). Each radial line represents 1 x 105 trajectories. (C) Probability density function of LIN-5 and dynein anaphase step-length distributions. (D) Determination of modeling parameters ρ, θfmax, θimax from Monte Carlo simulations and SSE calculations. (E) Determination of overlap for swivel model probability density functions for free swivel and interacting swivel 25th (0.028 μm) and 75th (0.0383 μm) percentiles, respectively. Green area represents overlap between theoretical 25th and 75th percentiles of free swivel and interacting swivel, respectively. (F) Probability density functions for posterior and anterior comparing best fit parameters of simulated data to experimental data.

We reasoned that the step length distributions for LIN-5 particles could be described by modeling G_α_—GPR1/2—LIN-5 as a rigid rod using the spherical coordinates (ρ,θ,φ), where ρ is the complex length, θ defines the polar deflection angle from the normal toward the membrane, and φ is the azimuthal angle about the base.

Experimentally measured LIN-5 trajectory turn angles were centered around 0° with a very broad spread (mean turn angle ± sd; 0.11° ± 88.4e°, ∼1.425 ∗ 10^7^ trajectories analyzed), consistent with a weak directional persistence and no preferred turning direction (Fig. 2C). The distribution indicated that LIN-5 trajectories contained a combination of random and directed turn angles. These results could reflect different subpopulations—short-timescale trajectories could display a turn angle randomness like a swivel undergoing rapid angular fluctuations, while long-timescale trajectories could reflect persistent, flow-aligned motion of the force generation complex with the cortex.

We modeled the azimuthal angle φ as uniformly distributed because dynein turn angles were uniformly distributed, and we expected LIN-5 to exhibit this behavior when engaged with a microtubule (as part of the force generation complex). To determine the maximum polar deflection angles, θ^fmax^ and θ^imax^, and the effective swivel length (ρ) of the G_α_—GPR1/2—LIN-5 complex, we performed “doppelgänger” Monte Carlo simulations over a range of candidate lengths (50 to 150 nm; (Garner *et al*., 2023)). For each ρ value, simulated step-length distributions were compared to experimental data using the sum of squared error (SSE) to identify best-fit parameters (Fig. 2D). Angular resolution improved steadily from ρ = 50 nm to 80 nm, plateaued between 80-90 nm, and then declined (Fig. 2D). The best-fit length (ρ = 80-90 nm) was within one pixel (length of 1 px) of our structural prediction (57 nm). The degree of agreement between simulated and experimental data exceeded that observed of dynein (simulated = 150 nm, predicted = 90 nm), likely reflecting the higher angular resolution achieved with LIN-5::mNG, which resides closer to the membrane (and the origin of TIRF’s evanescent field) than eGFP::DHC-1.

Using the best-fit value ρ = 90 nm, we identified parameterizations that minimized SSE and defined θ^fmax^ and θ^imax^. The angular resolution was highest at θ^fmax^ = [85°, 90°] for the free swivel group, and θ^imax^ = [40°, 45°] for the interacting swivel group (Fig. 2D). Step-length probability functions for each pole were generated by concatenating the corresponding free and interacting swivel simulations at these optimal angular parameters. The resulting theoretical distributions (Fig. 2E) provided a reasonable approximation of the experimental data for both anterior and posterior cortices, supporting the use of these parameter values for subsequent trajectory classifications.

Monte Carlo simulations were used to derive step-length probability mass functions for the free swivel and interacting swivel groups. The 25^th^ and 75^th^ percentiles of these distributions served as classification thresholds for assigning trajectories to each group. The 75th percentile of the interacting swivel distribution overlapped with the 25^th^ percentile of the free swivel distribution (Fig. 2F). To quantify the resulting uncertainty, we measured the proportion of overlapping steps and found that only 12% of interacting swivel step lengths overlapped with the free swivel distribution, and 16% of free swivel step-lengths overlapped with the interacting swivel distribution. The overlap percentages of the models indicated a reasonable level of confidence in using the 25^th^ and 75^th^ percentiles as thresholds for trajectory classification.

To validate our swivel model experimentally, we quantified the number of LIN-5 trajectories classified as interacting swivels in anterior and posterior cortices during anaphase—when posterior pulling forces position the mitotic spindle asymmetrically.

Trajectory counts were normalized to the cortical area analyzed for each embryo half. The density of interacting swivel trajectories was consistently higher in the posterior than the anterior cortex (mean ± SD: anterior = 70 ± 43 trajectories; posterior = 79 ± 24 trajectories; Fig. 3A), in agreement with our anaphase dynein data and previous reports. The ability to resolve this subtle yet reproducible difference at the single-molecule scale underscores the sensitivity of our modeling framework and highlights the precision with which cortical force asymmetry is established during the first division of the C. *elegans* zygote.

**Figure 3.**
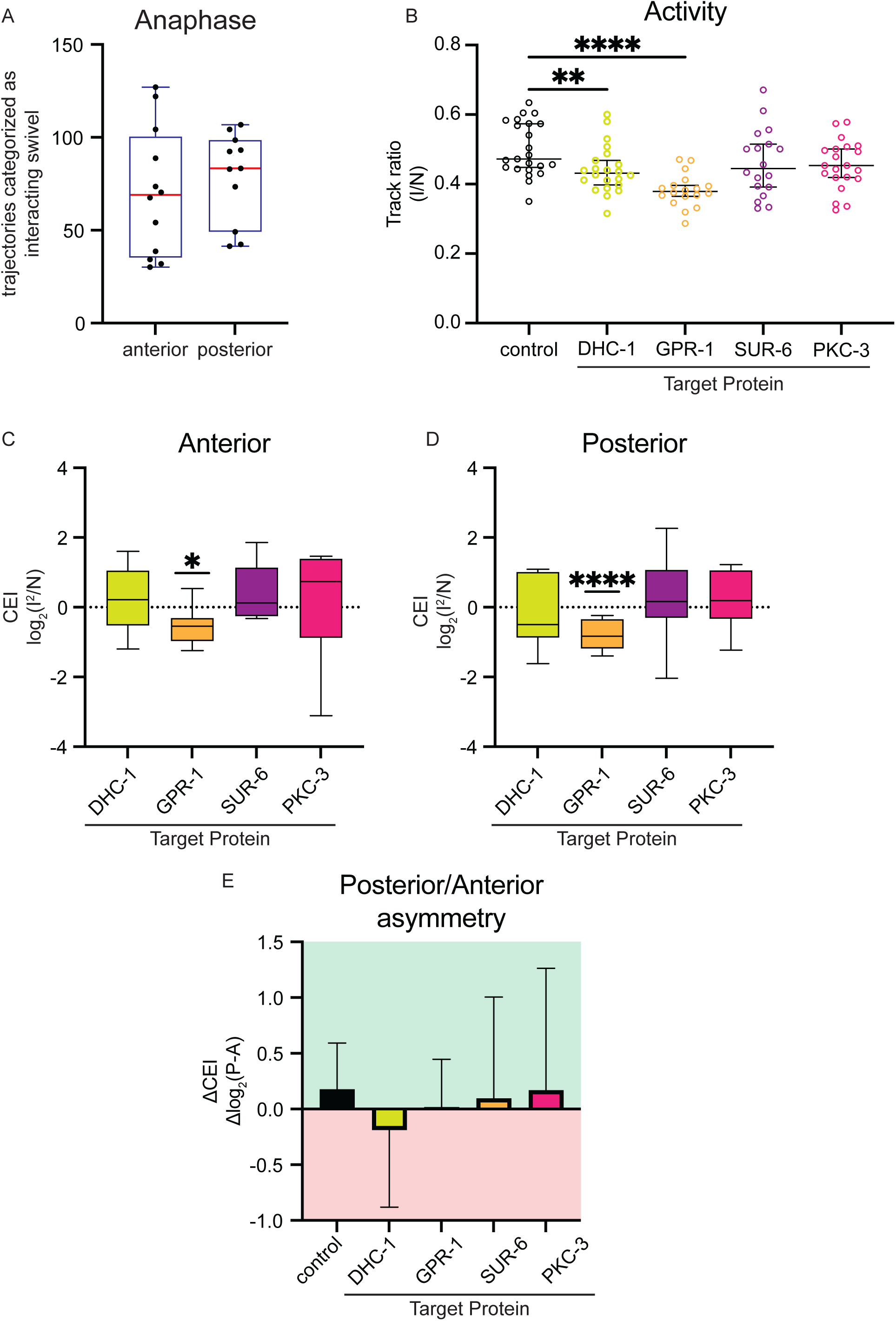
LIN-5 swivel model Activity and CEI metrics are sensitive to disruptions to the force generation complex hierarchy. (A) Total trajectories classified as swivel interacting in the anterior and posterior during anaphase in LIN-5::mNG embryos. The swivel model reproduced known force asymmetries during anaphase. (B) Activity metrics obtained from swivel data. Circles represent individual embryos. Black lines represent median ± 95% CI. (C–D) Baseline-subtracted box and whisker plots of CEI in the anterior (C) and posterior (D). Black lines represent 5th and 95th percentile. Statistics: Activity - Brown–Forsythe Anova and Welch’s t-test. CEI – one-way ANOVA and unpaired t-test. All t-tests for significance correspond to *p<0.05, **p<0.01, ***p<0.001, ****p<0.0001. (E) Posterior–anterior asymmetry measured as ΔCEI (posterior – anterior).

### Distinct contributions of DHC-1 and GPR-1 to LIN-5 cortical dynamics

We next examined how LIN-5 motion at the cortex is regulated by its key binding partners, dynein and GPR-1. GPR-1 links LIN-5 to the membrane, while dynein links LIN-5 and the rest of the force generation complex to the microtubule. We hypothesized depleting either linker would reduce LIN-5 Activity and CEI. Consistent with this prediction, Activity was significantly decreased in both *dhc-1(RNAi)* and *gpr-1(RNAi)* embryos, with the strongest reduction following GPR-1 depletion (Fig. 3B, Brown–Forsythe ANOVA, Welch’s t-test, *dhc-1(RNAi)* p = 0.0089, *gpr-1(RNAi)* p < 0.0001). CEI measurements revealed no significant change in *dhc-1(RNAi)* embryos relative to control in either cortical half, whereas *gpr-1(RNAi)* embryos showed significant decreases in both anterior and posterior cortices (Fig. 3C-D, one-way ANOVA, unpaired t-test anterior p = 0.0127; posterior p < 0.0001). However, ΔCEI was reversed in *dhc-1(RNAi)* embryos, while asymmetry was lost in *gpr-1(RNAi)* embryos (Fig. 3E). In all, these data indicate that DHC-1 and GPR-1 contribute to LIN-5 dynamics, albeit in different ways.

### PKC-3 and PP2A-B55/SUR-6 do not alter LIN-5 swivel dynamics

Having defined the swivel dynamics of LIN-5, we next investigated how its cortical behavior is modulated by phosphorylation and dephosphorylation. We hypothesized that SUR-6 regulates cortical microtubule interaction indirectly, for example by dephosphorylating LIN-5, the extensive phosphorylation of which is essential for its cortical function (Portegijs *et al*., 2016). The polarity kinase PKC-3 phosphorylates a subset of LIN-5, contributing to its asymmetric distribution (Portegijs *et al*., 2016). Despite these potential roles, neither *sur-6(RNAi)* nor *pkc-3(RNAi)* embryos exhibited major alterations in Activity or CEI (Fig. 3B–D). While *sur-6(RNAi)* and *pkc-3(RNAi)* embryos displayed greater variability in asymmetry, our analyses revealed no consistent deviation from control (Fig. 3E).

### LIN-5 is more abundant and longer-lived at the cortex than dynein

We next used our data to interpret the spatiotemporal relationship between LIN-5 and dynein (Movie S4. Like dynein, nearly all LIN-5 trajectories fell into either free swivel or interacting swivel categories, indicating the median step length of most trajectories fell between the 25^th^ and 75^th^ percentiles for free or interacting swivel groups (Linehan *et al*., 2023). However, compared to eGFP::dynein, LIN-5::mNG had a substantially longer cortical residence time (Brown–Forsythe ANOVA, Welch’s t-test; p < 0.0001). Average trajectory length for LIN-5::mNG particles during anaphase was 13.47 ± 1.99 steps in the anterior and 13.23 ± 1.273 steps in the posterior, whereas eGFP::DHC-1 particles traveled for averages of 8.16 ± 0.60 and 8.30 ± 0.21 steps, in the anterior and posterior respectively (Fig. 4A).

**Figure 4.**
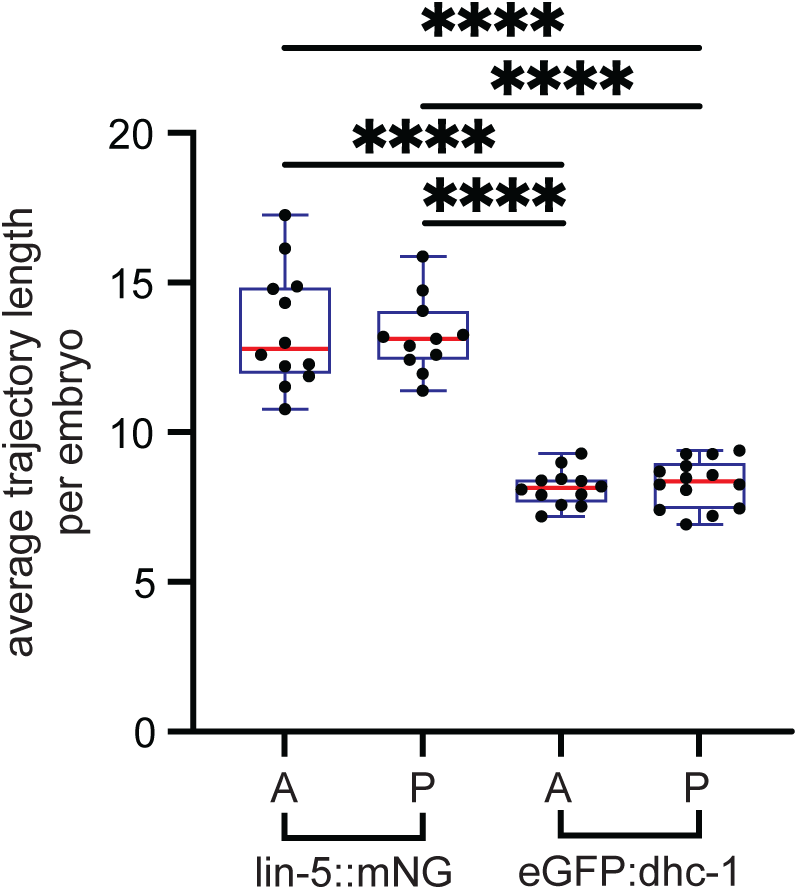
Comparing kinetics between LIN-5 and dynein. Box and whisker plots of average length in anterior and posterior during anaphase in lin-5::mNG and eGFP:dhc-1. Black dots represent individual embryos. A – anterior, P – posterior. Statistics: Brown–Forsythe ANOVA and Welch’s t-test. Welch’s t-tests for significance correspond to ****p<0.0001.

### LIN-5 dynamics couple short-term swivel motion with long-term cortical flows

We noticed that LIN-5 particles often moved in coordinated, network-like patterns, consistent with our earlier observation of a mild directional bias in turn-angle distribution (Fig. 2A). This suggested that LIN-5 motion operates on multiple time scales—rapid swivel-like fluctuations and slower, more persistent displacement.

To test this, we compared the behavior of short and long-duration trajectories by separating them at the average trajectory length threshold (13 steps per track). Short tracks (<13 steps) were predicted to behave as swivels with low directionality, whereas long tracks might display directed motion over time. As expected, longer trajectories exhibited greater displacement distances than shorter ones (Fig. 5A). Long (≥13 steps) trajectories flowed long distances (Fig. 5B and Supplemental Movie S5). This behavior contrasted with that of dynein, which consistently exhibited swivel behavior, and, rather, resembled previously reported cortical flows of GPR-1/2 that coincide in magnitude and direction with actomyosin cortical flows (De Simone *et al*., 2016).

**Figure 5.**
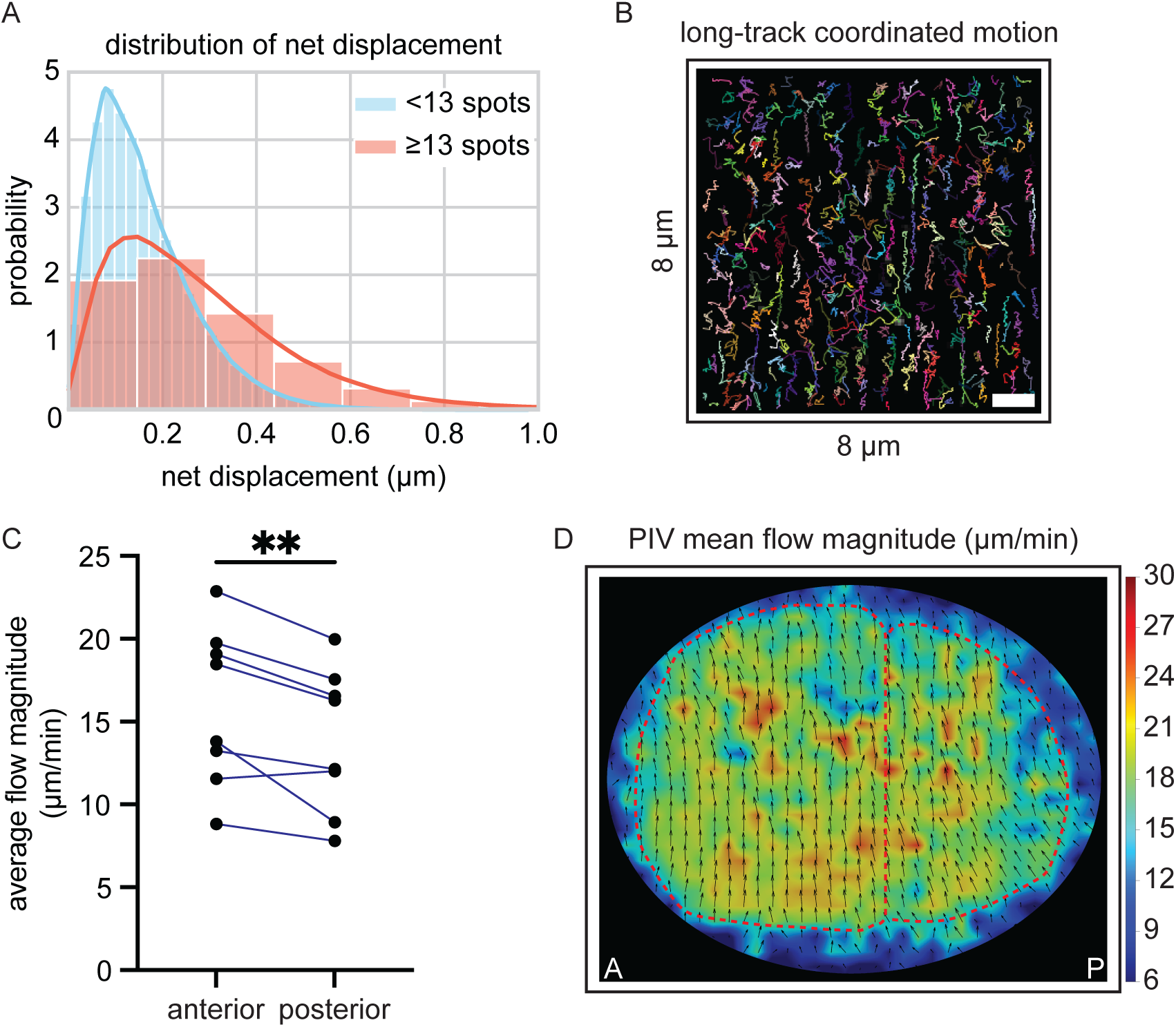
LIN-5 flows at long time scales during anaphase cortical rotation. (A) Probability density function of LIN-5 trajectories thresholded at median trajectory length (13 timepoints). (B) Overlay of longer tracks (>13 time points) from a representative image with single particle tracking. Note trajectories are directed along the y-axis. Scale bar = 1μm. (C) Comparison of average flow magnitude in anterior and posterior lin-5::mNG embryos. (D) Representative image of PIV analysis showing an overlay of mean flow magnitude. Relative magnitude is represented as a colormap. Arrows show direction of flow. Red dotted lines outline anterior and posterior halves. A - anterior, P – posterior. Statistics: Average flow magnitude – paired t-test. Paired t-test for significance corresponds to **p<0.01.

To quantify LIN-5’s coordinated behavior over time, we applied particle image velocimetry (PIV) to LIN-5::mNG embryos that displayed cortical rotation and anterior and posterior flows could be analyzed simultaneously (n = 8 embryos). Flow magnitudes were measured at one-twentieth of the acquisition rate (2 Hz). Mean flow magnitudes were comparable between the anterior (15.94 ± 4.70 μm min⁻¹) and posterior (13.91 ± 4.33 μm min⁻¹) halves, though consistently higher in the anterior (Fig. 5C–D; paired t-test, p = 0.0078). These flow velocities were within the range of previously reported actomyosin-derived cortical flows, suggesting that LIN-5 motion is influenced by cortical flow fields (Singh and Pohl, 2014; Middelkoop *et al*., 2021; Zaatri *et al*., 2021).

In contrast to dynein, LIN-5 displayed dynamic behaviors across two different time scales—rapid swivel-like motion at short intervals and bulk flow at longer ones. Together, these results reveal that LIN-5 integrates dynein-microtubule interactions at millisecond scale while collective flows emerge at the second scale.

## Discussion

During embryogenesis, the tightly regulated transmission of forces from the cortex through dynein and microtubules ensures proper centrosome and spindle positioning. These forces are mediated by the force generation complex, which transmits load from cortical dynein through LIN-5 and GPR-1/2 to the plasma membrane. Although the force generation complex’s components have been genetically defined, their dynamic behavior and regulation *in vivo* remained poorly understood. Using high-speed TIRF microscopy combined with quantitative single-particle analysis, we resolved the dynamics of dynein and LIN-5 at the cortex with unprecedented temporal precision. This approach, similar to recent single-molecule level HiLo microscopy of dynein in human cells describing dynein activation and microtubule capture during cargo transport (Tirumala *et al*., 2024), revealed distinct kinetic regimes of dynein and LIN-5 at the cortex, expanded our understanding of force generation complex regulation, and uncovered a previously unrecognized role for PP2A-B55/SUR-6 in shaping cortical force generation.

### PP2A-B55/SUR-6

Like many phosphatases, the mitotic regulator PP2A-B55/SUR-6 has traditionally been regarded as a housekeeping enzyme, acting primarily during the metaphase-to-anaphase transition to promote centrosome disassembly. However, recent work has broadened this limited view. Prior work demonstrated that SUR-6 cooperates with the nuclear lamina to promote centrosome separation, while SUR-6 regulates nuclear envelope breakdown through dephosphorylation of the lamin LMN-1 (Boudreau *et al*., 2019; Kapoor *et al*., 2023). Here, we uncovered a previously unrecognized role for PP2A-B55/SUR-6 in regulating cortical dynein–microtubule interactions. Both dynein Activity and CEI increased substantially following *sur-6(RNAi)* depletion, reflecting elevated total and interacting trajectories. This result implicates dynein as the mechanistic link through which SUR-6 influences cortical force production.

Because LIN-5, the cortical dynein-anchoring protein, is multi-phosphorylated and requires phosphorylation at specific residues for full functionality (Portegijs *et al*., 2016), we predicted that we would find evidence that LIN-5 is a direct substrate of SUR-6. However, LIN-5::mNG trajectory metrics remained unchanged upon SUR-6 depletion, suggesting that SUR-6 does not directly regulate LIN-5 Activity or CEI. Instead, SUR-6 may modulate dynein directly or modulate dynein localization and activity through an indirect mechanism.

Although dynein Activity at the cortex increased after SUR-6 depletion, SUR-6 enrichment could not be detected at the cortex in SUR-6::GFP (Bel Borja *et al*., 2020) and the apparent dynein accumulation at the male pronuclear envelope likely reflects secondary effects of nuclear size changes. Collectively, these findings define a role for PP2A-B55/SUR-6 in constraining dynein’s cortical engagement to maintain appropriate levels of force generation.

PAR proteins spatially restrict GPR-1/2 and thereby LIN-5 localization (Goldstein and Macara, 2007). Our results raise the possibility that phosphoregulation by SUR-6 acts in parallel with PAR-dependent pathways to modulate cortical force asymmetry. PP2A-mediated dephosphorylation could fine-tune dynein’s cortical residence or microtubule-binding capacity in a spatially restricted manner, allowing mechanical feedback between polarity and force generation machinery.

### LIN-5: Swivel and Flow

Here, we provided the first detailed description of LIN-5 dynamics at the cortex. We quantified LIN-5 trajectories and found striking kinetic distinctions between LIN-5 and dynein. LIN-5 exhibited significantly longer residence times than dynein, reflecting LIN-5’s stable association with the membrane (Fig. 4). Despite these differences, LIN-5 behavior could be captured using our swivel model classification framework, originally designed for dynein. This model accurately estimated the effective length of the force generation complex containing LIN-5::mNG (Fig. 2D), recapitulated known anaphase force asymmetries (Fig. 3A), and remained sensitive to reductions in force generation complex efficacy via *dhc-1(RNAi)* and in cortical anchoring via *gpr-1(RNAi)* (Fig. 3B–E).

At short time scales, LIN-5 behaved similarly to dynein, exhibiting rapid swivel-like fluctuations consistent with transient microtubule engagement (Fig. 2A). Over longer intervals, however, LIN-5’s extended cortical residence resulted in coherent, flow-like motion across the cortex (Fig. 5). These flows occurred on the same order of magnitude of actomyosin flows during anaphase cortical rotation reported by others (Singh and Pohl, 2014; Middelkoop *et al*., 2021; Zaatri *et al*., 2021).

Our findings suggest a kinetic division of labor within the force generation complex, where LIN-5 provides spatial persistence at the membrane while dynein generates transient, load-bearing interactions. This dynamic complementarity may enable rapid modulation of cortical forces without compromising positional stability, a feature shared with NuMA–dynein complexes in vertebrate cells (Hueschen *et al*., 2019). Viewed as a modular system, the force generation complex integrates motors, adaptors, and regulatory enzymes that tune cortical forces across space and time. Dynein determines instantaneous pulling events, LIN-5 establishes cortical persistence, and phosphoregulators such as SUR-6 adjust the balance between these modes to maintain overall mechanical symmetry. While we did not observe dynein flows with our imaging scheme, we cannot exclude the possibility that dynein flows occurred just outside the imaging depth. It will also be of importance to directly measure interactions between the force generation complex and the actomyosin cortex simultaneously, which may provide more evidence for coupling between the microtubule and actin cytoskeletons.

By linking molecular kinetics to emergent cortical behavior, our study bridges the gap between single-molecule dynamics and cell-scale force generation. The conceptual and analytical framework developed here provides a foundation for exploring how dynamic cytoskeletal assemblies achieve mechanical coordination across space and time in diverse morphogenetic contexts.

## Acknowledgements

We greatly thank Tony Perdue and Nathanaël Prunet (Microscopy Core, Biology Department, UNC-Chapel Hill) for their support. We thank the Bob Goldstein Lab for strains and reagents. We also thank Kacy Gordon and Stephanie Gupton for critical reading and discussion of the manuscript. This study was supported by the National Science Foundation CAREER Award 1652512 to P.S.M. J.B.L. was supported in part by a grant from NIGMS under award T32 GM119999. This study was also supported by the NIGMS of the National Institutes of Health (R35GM144238) and by the National Science Foundation (2153790) to A.S.M.

## Conflict of Interest

The authors declare no competing financial interests.

## Supplemental Movie Legend

**Movie S1. Time-lapse demonstrating single particle tracking of SV1803 with image (left) and tracking (right) separated for visibility.** Scale bar = 1 μm. Real time (40 Hz). Related to Fig. 1.

**Movie S2. Time-lapse comparing “Raw” vs “DoG” filtered images of LP585.** Scale bar = 1 μm. Real time (40 Hz). Related to Fig. S2.

**Movie S3. Time-lapse demonstrating SPT of LP585 with tracks (random color) overlaid on image.** Scale bar = 1 μm. Real time (40 Hz). Related to Fig. 3.

**Movie S4. Time-lapse comparing SV1803 and LP585 during anaphase**. Combined movies S1 and S3 (“Raw”). Real time (40 Hz). Related to Fig. 4.

**Movie S5. Time-lapse demonstrating LP585 flows.** Movie is 5x real time (200 Hz). Related to Fig. 5.

---

**Figure 1 – figure supplement S1.**
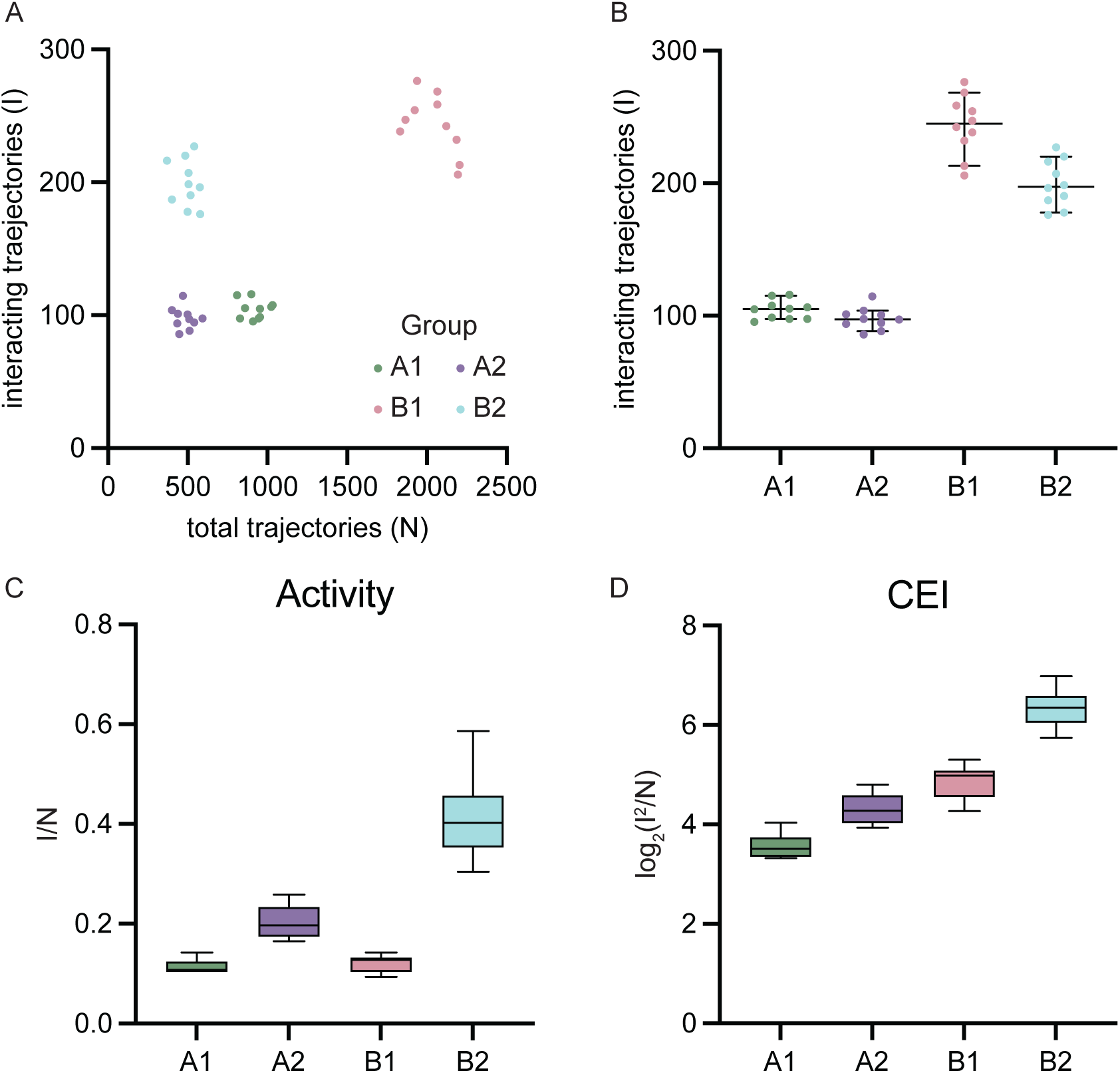
In silico comparison for four hypothetical groups demonstrates how Activity and collective efficiency index (CEI) measurements enable the comparison of the relationship between cortical dynein number and dynein-microtubule interactions. Four populations were generated where interacting/total trajectories means of the four groups are: A1–100/1000; A2–100/500; B1–250/2000;B2–250/500. 10 replicates were produced for each group with 10% random relative noise distributed normally about the mean. (A) Scatter plot of four hypothetical groups comparing total trajectories to interacting trajectories. The four groups are spatially distinct. (B) Plot of interacting trajectories for each group. Black lines indicate median with 95% CI. Dots represent replicates. (C–D) Box and whisker plots of Activity (C) and collective efficiency index (D) for four hypothetical groups. Note the complimentary differences in C–D that are lost by considering only B.

**Figure 2 – figure supplement S2.**
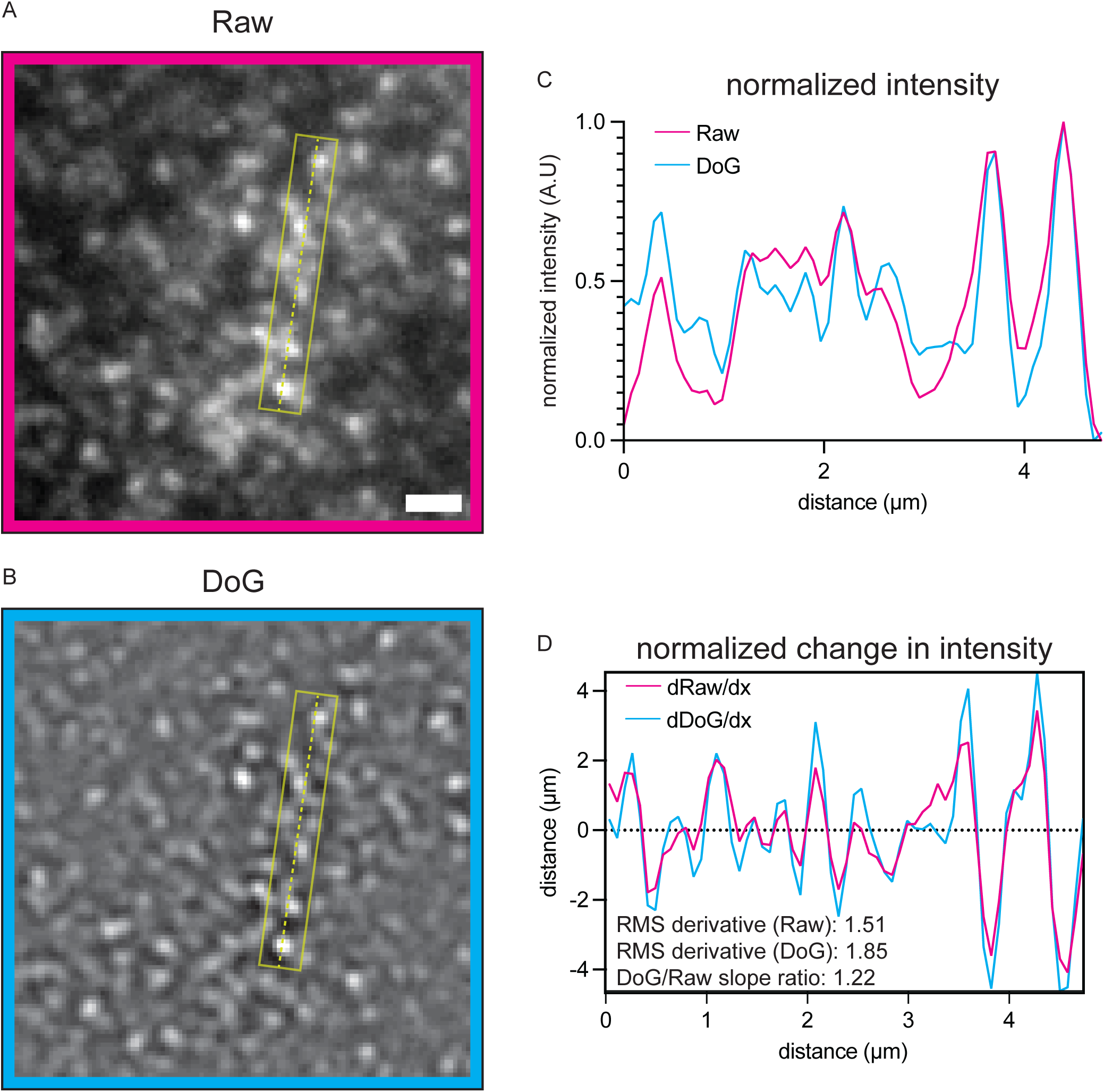
Difference of Gaussian (DoG) filtering improves spot detection. (A–B) “Raw” (A) and “DoG” processed (σ=1.5-1) (B) TIRF images of LP585. Dotted yellow line represents line scan used for measuring intensity values (direction–top to bottom) (A.U.). Yellow rectangle indicates the area around the line scan included to measure intensity values. (C–D) Normalized intensity (C) and normalized change in intensity (dy/dx) for “Raw” and “Dog” images. Note the improved contrast of “DoG” in (D) (cyan line maximum and minimum values increased, and more frequent sign change occurrence (dotted line at 0).

## Notes

### Competing Interest Statement

The authors have declared no competing interest.

## References

Barisic, M, Aguiar, P, Geley, S, and Maiato, H (2014). Kinetochore motors drive congression of peripheral polar chromosomes by overcoming random arm-ejection forces. Nat Cell Biol 16, 1249–1256.

Bel Borja, L, Soubigou, F, Taylor, SJP, Fraguas Bringas, C, Budrewicz, J, Lara-Gonzalez, P, Sorensen Turpin, CG, Bembenek, JN, Cheerambathur, DK, and Pelisch, F (2020). BUB-1 targets PP2A:B56 to regulate chromosome congression during meiosis I in C. elegans oocytes. ELife 9.

Boudreau, V, Chen, R, Edwards, A, Sulaimain, M, and Maddox, PS (2019). PP2A-B55/SUR-6 collaborates with the nuclear lamina for centrosome separation during mitotic entry. Mol Biol Cell 30, 876–886.

Crisp, M, Liu, Q, Roux, K, Rattner, JB, Shanahan, C, Burke, B, Stahl, PD, and Hodzic, D (2006). Coupling of the nucleus and cytoplasm: role of the LINC complex. J Cell Biol 172, 41–53.

De Simone, A, and Gönczy, P (2017). Computer simulations reveal mechanisms that organize nuclear dynein forces to separate centrosomes. Mol Biol Cell 28, 3165–3170.

De Simone, A, Nédélec, F, and Gönczy, P (2016). Dynein transmits polarized actomyosin cortical flows to promote centrosome separation. Cell Rep 14, 2250–2262.

De Simone, A, Spahr, A, Busso, C, and Gönczy, P (2018). Uncovering the balance of forces driving microtubule aster migration in C. elegans zygotes. Nat Commun 9, 938.

Ershov, D, Phan, M-S, Pylvänäinen, JW, Rigaud, SU, Le Blanc, L, Charles-Orszag, A, Conway, JR, Laine, RF, Roy, NH, and Bonazzi, D (2021). Bringing TrackMate into the era of machine-learning and deep-learning. BioRxiv, 2021–09.

Ershov, D et al. (2022). TrackMate 7: integrating state-of-the-art segmentation algorithms into tracking pipelines. Nat Methods 19, 829–832.

Fielmich, L-E, Schmidt, R, Dickinson, DJ, Goldstein, B, Akhmanova, A, and van den Heuvel, S (2018). Optogenetic dissection of mitotic spindle positioning in vivo. ELife 7.

Fraser, AG, Kamath, RS, Zipperlen, P, Martinez-Campos, M, Sohrmann, M, and Ahringer, J (2000). Functional genomic analysis of C. elegans chromosome I by systematic RNA interference. Nature 408, 325–330.

Garner, RM, Molines, AT, Theriot, JA, and Chang, F (2023). Vast heterogeneity in cytoplasmic diffusion rates revealed by nanorheology and Doppelgänger simulations. Biophys J 122, 767–783.

Goldstein, B, and Macara, IG (2007). The PAR proteins: fundamental players in animal cell polarization. Dev Cell 13, 609–622.

Gönczy, P, Pichler, S, Kirkham, M, and Hyman, AA (1999). Cytoplasmic dynein is required for distinct aspects of MTOC positioning, including centrosome separation, in the one cell stage Caenorhabditis elegans embryo. J Cell Biol 147, 135–150.

Gusnowski, EM, and Srayko, M (2011). Visualization of dynein-dependent microtubule gliding at the cell cortex: implications for spindle positioning. J Cell Biol 194, 377–386.

Heppert, JK, Pani, AM, Roberts, AM, Dickinson, DJ, and Goldstein, B (2018). A CRISPR Tagging-Based Screen Reveals Localized Players in Wnt-Directed Asymmetric Cell Division. Genetics 208, 1147–1164.

Huang, J, Roberts, AJ, Leschziner, AE, and Reck-Peterson, SL (2012). Lis1 acts as a “clutch” between the ATPase and microtubule-binding domains of the dynein motor. Cell 150, 975–986.

Hueschen, CL, Galstyan, V, Amouzgar, M, Phillips, R, and Dumont, S (2019). Microtubule End-Clustering Maintains a Steady-State Spindle Shape. Curr Biol 29, 700–708.e5.

Jankele, R, Jelier, R, and Gönczy, P (2021). Physically asymmetric division of the C. elegans zygote ensures invariably successful embryogenesis. ELife 10.

Jumper, J et al. (2021). Highly accurate protein structure prediction with AlphaFold. Nature 596, 583–589.

Kamath, RS, Martinez-Campos, M, Zipperlen, P, Fraser, AG, and Ahringer, J (2001). Effectiveness of specific RNA-mediated interference through ingested double-stranded RNA in Caenorhabditis elegans. Genome Biol 2, RESEARCH0002.

Kapoor, S, Adhikary, K, and Kotak, S (2023). PP2A-B55SUR-6 promotes nuclear envelope breakdown in C. elegans embryos. Cell Rep 42, 113495.

Kiyomitsu, T (2019). The cortical force-generating machinery: how cortical spindle-pulling forces are generated. Curr Opin Cell Biol 60, 1–8.

Laan, L, Pavin, N, Husson, J, Romet-Lemonne, G, van Duijn, M, López, MP, Vale, RD, Jülicher, F, Reck-Peterson, SL, and Dogterom, M (2012a). Cortical dynein controls microtubule dynamics to generate pulling forces that position microtubule asters. Cell 148, 502–514.

Laan, L, Roth, S, and Dogterom, M (2012b). End-on microtubule-dynein interactions and pulling-based positioning of microtubule organizing centers. Cell Cycle 11, 3750–3757.

Linehan, JB, Edwards, GA, Boudreau, V, Maddox, AS, and Maddox, PS (2023). Model-based trajectory classification of anchored molecular motor-biopolymer interactions. Biophysical Reports 3, 100130.

Mazel, T, Biesemann, A, Krejczy, M, Nowald, J, Müller, O, and Dehmelt, L (2014). Direct observation of microtubule pushing by cortical dynein in living cells. Mol Biol Cell 25, 95–106.

Merdes, A, Ramyar, K, Vechio, JD, and Cleveland, DW (1996). A complex of NuMA and cytoplasmic dynein is essential for mitotic spindle assembly. Cell 87, 447–458.

Middelkoop, TC, Garcia-Baucells, J, Quintero-Cadena, P, Pimpale, LG, Yazdi, S, Sternberg, PW, Gross, P, and Grill, SW (2021). CYK-1/Formin activation in cortical RhoA signaling centers promotes organismal left-right symmetry breaking. Proc Natl Acad Sci USA 118.

Miura, K (2020). Bleach correction ImageJ plugin for compensating the photobleaching of time-lapse sequences. F1000Res 9, 1494.

O’Rourke, SM, Christensen, SN, and Bowerman, B (2010). Caenorhabditis elegans EFA-6 limits microtubule growth at the cell cortex. Nat Cell Biol 12, 1235–1241.

Okumura, M, Natsume, T, Kanemaki, MT, and Kiyomitsu, T (2018). Dynein-Dynactin-NuMA clusters generate cortical spindle-pulling forces as a multi-arm ensemble. ELife 7.

Park, DH, and Rose, LS (2008). Dynamic localization of LIN-5 and GPR-1/2 to cortical force generation domains during spindle positioning. Dev Biol 315, 42–54.

Portegijs, V, Fielmich, L-E, Galli, M, Schmidt, R, Muñoz, J, van Mourik, T, Akhmanova, A, Heck, AJR, Boxem, M, and van den Heuvel, S (2016). Multisite Phosphorylation of NuMA-Related LIN-5 Controls Mitotic Spindle Positioning in C. elegans. PLoS Genet 12, e1006291.

Roberts, AJ, Kon, T, Knight, PJ, Sutoh, K, and Burgess, SA (2013). Functions and mechanics of dynein motor proteins. Nat Rev Mol Cell Biol 14, 713–726.

Robinson, JT, Wojcik, EJ, Sanders, MA, McGrail, M, and Hays, TS (1999). Cytoplasmic dynein is required for the nuclear attachment and migration of centrosomes during mitosis in Drosophila. J Cell Biol 146, 597–608.

Rodriguez-Garcia, R, Chesneau, L, Pastezeur, S, Roul, J, Tramier, M, and Pécréaux, J (2018). The polarity-induced force imbalance in Caenorhabditis elegans embryos is caused by asymmetric binding rates of dynein to the cortex. Mol Biol Cell 29, 3093–3104.

Salina, D, Bodoor, K, Eckley, DM, Schroer, TA, Rattner, JB, and Burke, B (2002). Cytoplasmic dynein as a facilitator of nuclear envelope breakdown. Cell 108, 97–107.

Schindelin, J et al. (2012). Fiji: an open-source platform for biological-image analysis. Nat Methods 9, 676–682.

Schmidt, R, Fielmich, L-E, Grigoriev, I, Katrukha, EA, Akhmanova, A, and van den Heuvel, S (2017). Two populations of cytoplasmic dynein contribute to spindle positioning in C. elegans embryos. J Cell Biol 216, 2777–2793.

Singh, D, and Pohl, C (2014). Coupling of rotational cortical flow, asymmetric midbody positioning, and spindle rotation mediates dorsoventral axis formation in C. elegans. Dev Cell 28, 253–267.

Splinter, D et al. (2010). Bicaudal D2, dynein, and kinesin-1 associate with nuclear pore complexes and regulate centrosome and nuclear positioning during mitotic entry. PLoS Biol 8, e1000350.

Thielicke, W, and Sonntag, R (2021). Particle Image Velocimetry for MATLAB: Accuracy and enhanced algorithms in PIVlab. J Open Res Softw 9.

Thielicke, W (2022). Pulse-length induced motion blur in PIV particle images: To be avoided at any cost?

Tinevez, J-Y, Perry, N, Schindelin, J, Hoopes, GM, Reynolds, GD, Laplantine, E, Bednarek, SY, Shorte, SL, and Eliceiri, KW (2017). TrackMate: An open and extensible platform for single-particle tracking. Methods 115, 80–90.

Tirumala, NA, Redpath, GMI, Skerhut, SV, Dolai, P, Kapoor-Kaushik, N, Ariotti, N, Vijay Kumar, K, and Ananthanarayanan, V (2024). Single-molecule imaging of stochastic interactions that drive dynein activation and cargo movement in cells. J Cell Biol 223.

Varadi, M et al. (2022). AlphaFold Protein Structure Database: massively expanding the structural coverage of protein-sequence space with high-accuracy models. Nucleic Acids Res 50, D439–D444.

Vorozhko, VV, Emanuele, MJ, Kallio, MJ, Stukenberg, PT, and Gorbsky, GJ (2008). Multiple mechanisms of chromosome movement in vertebrate cells mediated through the Ndc80 complex and dynein/dynactin. Chromosoma 117, 169–179.

Yang, Z, Tulu, US, Wadsworth, P, and Rieder, CL (2007). Kinetochore dynein is required for chromosome motion and congression independent of the spindle checkpoint. Curr Biol 17, 973–980.

Zaatri, A, Perry, JA, and Maddox, AS (2021). Septins and a formin have distinct functions in anaphase chiral cortical rotation in the Caenorhabditis elegans zygote. Mol Biol Cell 32, 1283–1292.

Zou, S, Toropova, K, Roberts, AJ, Reck-Peterson, SL, and Leschziner, AE (2014). Lis1 regulates dynein as a molecular wedge. Biophys J 106, 353a.

